# Spatial Mechanomics for Tissue-Scale Biomechanical Mapping and Multi-omics Integration

**DOI:** 10.64898/2026.02.25.703280

**Authors:** Weichang Xie, Zichen Wang, Qiji Shan, Qiang Zhao, Xiaofeng Ye

## Abstract

Tissue mechanical properties are spatially heterogeneous and tightly coupled to cellular function, developmental patterning, and disease progression, yet spatially resolved characterization of viscoelastic and microrheological behavior across intact tissues remains limited. Here we introduce spatial mechanomics, a framework for tissue-wide acquisition, quantitative extraction, and computational representation of location-resolved mechanical states. Using BioAFM-based spatial sampling with multi-protocol microrheology, we acquire force responses at defined tissue coordinates and fit physically interpretable viscoelastic models to extract elastic, viscous, and frequency-dependent parameters at each position. These parameters are assembled into per-niche mechanomic feature vectors and reconstructed into tissue-scale mechanomic atlases that resolve heterogeneous mechanical organization. We implement these capabilities in MechScape, an open-source computational platform that supports force curve fitting, spatial feature matrix construction, unsupervised domain discovery, and cross-modal alignment with histological and molecular measurements. Application to murine myocardial tissue reveals that spatial mechanomics identifies distinct mechanical states, quantifies condition-dependent remodeling across all measured parameters, and resolves spatially coherent mechanical domains. This work establishes spatial mechanomics as a quantitative approach for tissue-scale biomechanical mapping and provides a generalizable framework for integrating mechanics as an omics layer in multi-modal tissue analysis.

## Introduction

The mechanical properties of biological tissues are not merely passive structural attributes but active regulators of cellular behavior, tissue homeostasis, and disease progression^1–3^. Cells continuously sense and respond to the stiffness, viscoelasticity, and topography of their microenvironment through mechanotransduction pathways that influence proliferation, differentiation, migration, and apoptosis^4,5^. In pathological contexts, altered tissue mechanics both reflect and drive disease: fibrotic tissues stiffen through excessive extracellular matrix deposition, tumors exhibit characteristic mechanical heterogeneity, and cardiovascular diseases fundamentally remodel the viscoelastic properties of myocardial tissue^6–8^. Understanding how mechanical properties are spatially organized within tissues and how this organization changes in disease is therefore central to both basic biology and translational medicine.

Despite the recognized importance of tissue mechanics, current measurement approaches face fundamental limitations in spatial resolution and parametric breadth. Bulk rheology and conventional mechanical testing provide tissue-averaged properties but sacrifice spatial information entirely^9^. Atomic force microscopy (AFM) has emerged as a powerful tool for nanoscale and microscale mechanical characterization, enabling measurement of local elastic moduli, adhesion forces, and viscoelastic properties^10,11^. However, most AFM-based studies measure a single mechanical parameter—typically Young’s modulus—at isolated positions, missing the rich, multi-parametric mechanical landscape that defines tissue mechanical identity. Moreover, the analytical pipeline from raw force curves to spatially mapped mechanical features has not been systematized in a manner analogous to the computational frameworks that have transformed genomics, transcriptomics, and proteomics into scalable, integrative omics disciplines^12–14^.

The concept of an omics-scale mechanical characterization—which we term *spatial mechanomics*— requires advances along three axes. First, measurement protocols must capture multiple complementary mechanical responses at each tissue position, including quasi-static elasticity, time-dependent viscoelastic relaxation, adhesion, and frequency-dependent dynamic moduli. Second, physically interpretable constitutive models must be fitted to these responses to extract quantitative parameters that are comparable across conditions and amenable to downstream analysis. Third, computational infrastructure must support assembly of these parameters with other omics modalities to enable mechanistic interpretation.

Here we present a spatial mechanomics framework that addresses these requirements (Fig. 1). We develop a multi-protocol BioAFM sampling strategy that acquires location-resolved force spectroscopy comprising approach–retract curves, constant-height creep, constant-force relaxation, and multi-frequency oscillatory microrheology within a single measurement cycle at each tissue position. Systematic models fitting to extract mechanical parameters per niche, including spanning elasticity, adhesion, viscoelastic time constants, and frequency-dependent storage and loss moduli are implemented. These per-niche feature vectors are reconstructed into spatial mechanomic maps that reveal tissue-scale mechanical organization. The complete analytical pipeline is implemented in MechScape, an open-source software platform that serves as an integrative toolkit for spatial mechanomics analysis. We validate this framework on murine cardiac tissue comparing sham and myocardial infarction conditions, demonstrating that spatial mechanomics resolves condition-dependent mechanical remodeling, identifies spatially coherent mechanical domains through unsupervised analysis.

**Fig. 1.**
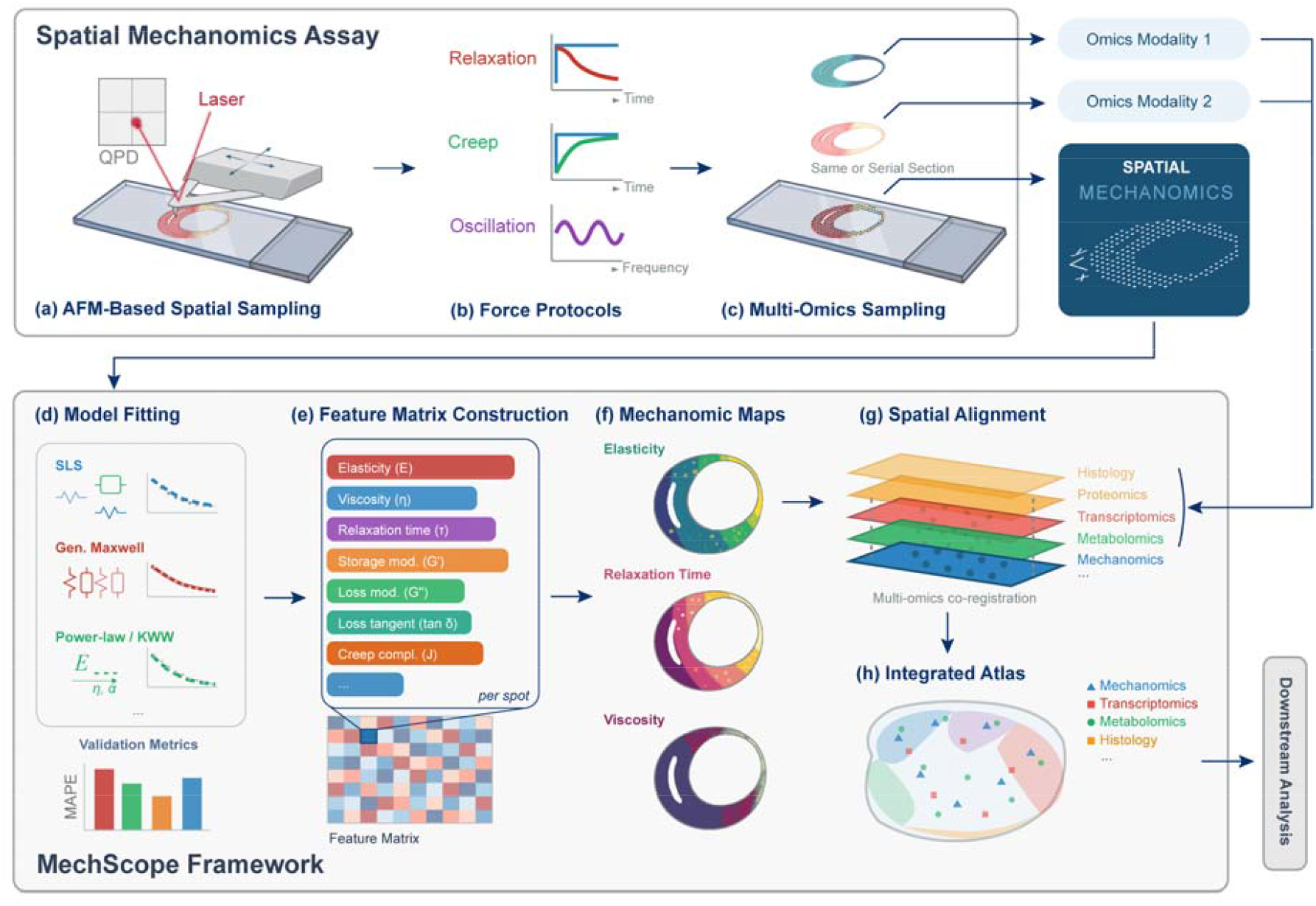
Spatial mechanomics enables tissue-scale microrheological mapping and integrative analysis. (a) BioAFM-based spatial sampling acquires location-resolved mechanical responses across intact tissue. (b) Multi-protocol force spectroscopy captures complementary viscoelastic and microrheological dynamics at each spatial position. (c) Spatially matched sections enable cross-modal measurements. (d–e) Physically interpretable viscoelastic models are fitted to force responses to generate mechanomic feature representations. (f) Reconstructed parameter fields form tissue-scale mechanomic maps. (g–h) Spatial co-registration integrates mechanomic data with histological and molecular modalities to generate an integrated atlas for downstream analysis.

## Results

### Spatial mechanomics framework for tissue-scale mechanical profiling

We designed a spatial mechanomics workflow that transforms raw BioAFM force measurements into tissue-scale, multi-parametric mechanical atlases (Fig. 1). The framework operates through four stages: spatially registered force acquisition, model-based parameter extraction, feature matrix assembly, and integrative analysis. At the acquisition stage, intact cryosectioned tissue is mounted on a BioAFM platform and a multi-protocol force spectroscopy routine is executed at each position of a predefined spatial grid (Fig. 1a). Each measurement cycle captures a force–indentation approach curve, a constant-height creep segment, a constant-force relaxation segment, multi-frequency sinusoidal oscillation, and a retraction curve, yielding complementary readouts of elastic, viscous, and dynamic mechanical behavior from a single spatial coordinate (Fig. 1b).

Serial tissue sections enable spatially matched measurements across modalities, allowing co-registration of mechanomic data with other omics on same or adjacent sections (Fig. 1c). At each sampled position, raw force responses are fitted with constitutive viscoelastic models selected via information-theoretic criteria to extract a quantitative parameter vector encompassing Young’s modulus, adhesion force, creep and relaxation time constants, and frequency-dependent storage and loss moduli. These per-niche feature vectors are assembled into a spatial feature matrix—the mechanomic representation of the tissue (Fig. 1d– f). Downstream analysis includes spatial reconstruction of parameter fields to generate mechanomic maps, unsupervised dimensionality reduction and clustering to identify mechanical domains, and cross-modal alignment with histological and molecular data to construct integrated tissue atlases (Fig. 1g–h).

### Model-based extraction of microrheological parameters from multi-protocol force spectroscopy

To extract quantitative mechanical parameters from each force spectroscopy cycle, we developed a systematic model fitting and selection pipeline (Fig. 2). A programmable 10-segment force protocol was executed at each spatial position using a BioAFM with a spherical-tipped cantilever. The protocol comprises an initial approach to contact, a constant-height creep hold, a brief retraction step, a force-triggered transition, a constant-force clamp, sinusoidal oscillation at four frequencies (1, 10, 100, and 500 Hz), and a final retraction (Fig. 2a). This multi-protocol design captures complementary viscoelastic information—time-domain creep and relaxation dynamics as well as frequency-domain dynamic moduli—within a single measurement cycle at each tissue position.

**Fig. 2.**
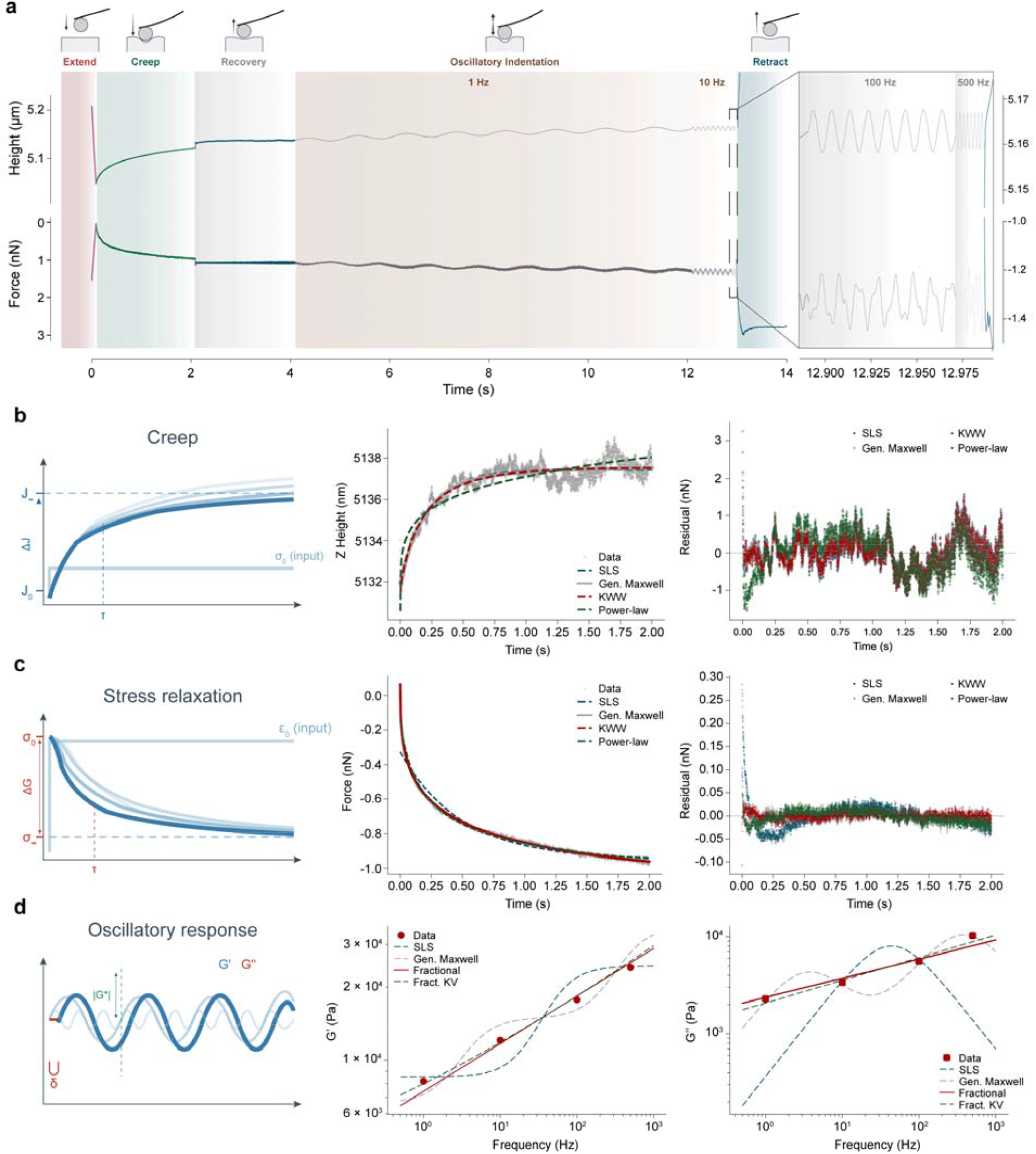
Model-based extraction of microrheological parameters from force spectroscopy. (a) Representative force protocols combining extension, creep, recovery, oscillatory indentation, and retraction within a single measurement cycle. Height and force responses reflect time- and frequency-dependent mechanical behavior. (b) Creep responses under constant load with representative fits using physically interpretable viscoelastic models and corresponding residual analysis. (c) Stress relaxation responses under step deformation with model fitting and residual evaluation. (d) Oscillatory microrheology across frequencies yields storage and loss moduli, with model-based reconstruction of frequency-dependent mechanical spectra.

Under constant-force feedback, height recovery reflecting creep compliance was recorded and fitted with four constitutive models: the standard linear solid (SLS/Zener, 3 parameters), a two-arm generalized Maxwell model (5 parameters), the Kohlrausch–Williams–Watts stretched exponential (KWW, 4 parameters), and a power-law model (4 parameters) (Fig. 2b). Model selection was performed using the corrected Akaike Information Criterion (AICc). The two-arm generalized Maxwell model achieved the best AICc (R^2^ = 0.8856), closely followed by SLS (R^2^ = 0.8835) and KWW (R^2^ = 0.8818). Lower overall R^2^ values compared with the creep segment reflect the intrinsic noise of feedback-controlled height measurements. Instantaneous and equilibrium elastic moduli (E□, E∞) were derived from the fitted force values using Hertz contact mechanics for a spherical indenter.

Force relaxation during the constant-height creep segment was fitted with analogous viscoelastic models (Fig. 2c). The KWW model provided the best fit (ΔAIC = 0; R^2^ = 0.9974), yielding an equilibrium force F∞ = −1.173 ± 0.004 nN, a characteristic time τ = 0.456 ± 0.005 s, and a stretched exponent β = 0.387 ± 0.003. The stretched exponent β < 1 indicates a broad distribution of relaxation times, characteristic of structurally heterogeneous viscoelastic media. The generalized Maxwell model ranked second (R^2^ = 0.9968; ΔAIC = 815), resolving fast (τ1 = 0.095 s) and slow (τ2 = 0.997 s) relaxation modes, while the single-exponential SLS (R^2^ = 0.9684) and power-law (R^2^ = 0.9929) models were substantially inferior.

Frequency-dependent viscoelastic moduli were extracted from oscillatory segments by Fourier analysis of force and displacement signals (Fig. 2d). Storage modulus G′ and loss modulus G′′ were computed from the amplitude ratio and phase lag at each drive frequency. Both moduli increased monotonically over the 1–500 Hz range, from G′ = 8,177 Pa and G′′ = 2,296 Pa at 1 Hz to G′ = 24,157 Pa and G′′ = 10,266 Pa at 500 Hz. G′ exceeded G′′ at all frequencies (tan δ < 0.5), confirming a predominantly elastic response with increasing viscous dissipation at higher frequencies (tan δ rising from 0.28 to 0.43). Simultaneous fitting of the frequency-dependent spectra selected a fractional power-law model (R^2^ = 0.981), yielding G□ = 5,436 ± 520 Pa·s and a power-law exponent α = 0.197 ± 0.012. This low exponent is consistent with soft glassy or cytoskeletal-like rheology, where α = 0 corresponds to a purely elastic solid and α = 0.5 to a Rouse-like polymer solution. These results demonstrate that the multi-protocol measurement and model selection pipeline extracts physically meaningful, multi-scale mechanical parameters suitable for constructing mechanomic feature representations.

### Spatial mechanomic atlases resolve tissue-scale mechanical remodeling in cardiac disease

To evaluate whether spatial mechanomics can resolve condition-dependent mechanical organization, we applied the framework to murine cardiac tissue comparing Sham and myocardial infarction (MI) samples. Tissue sections were cryosectioned, spatially sampled by BioAFM, and serially stained with Masson’s trichrome to enable histological co-registration (Fig. 3a). At each niche, the full 10-segment force protocol was executed, and 20 mechanical parameters were extracted per position, generating spatially resolved mechanomic maps for each parameter across both conditions.

**Fig. 3.**
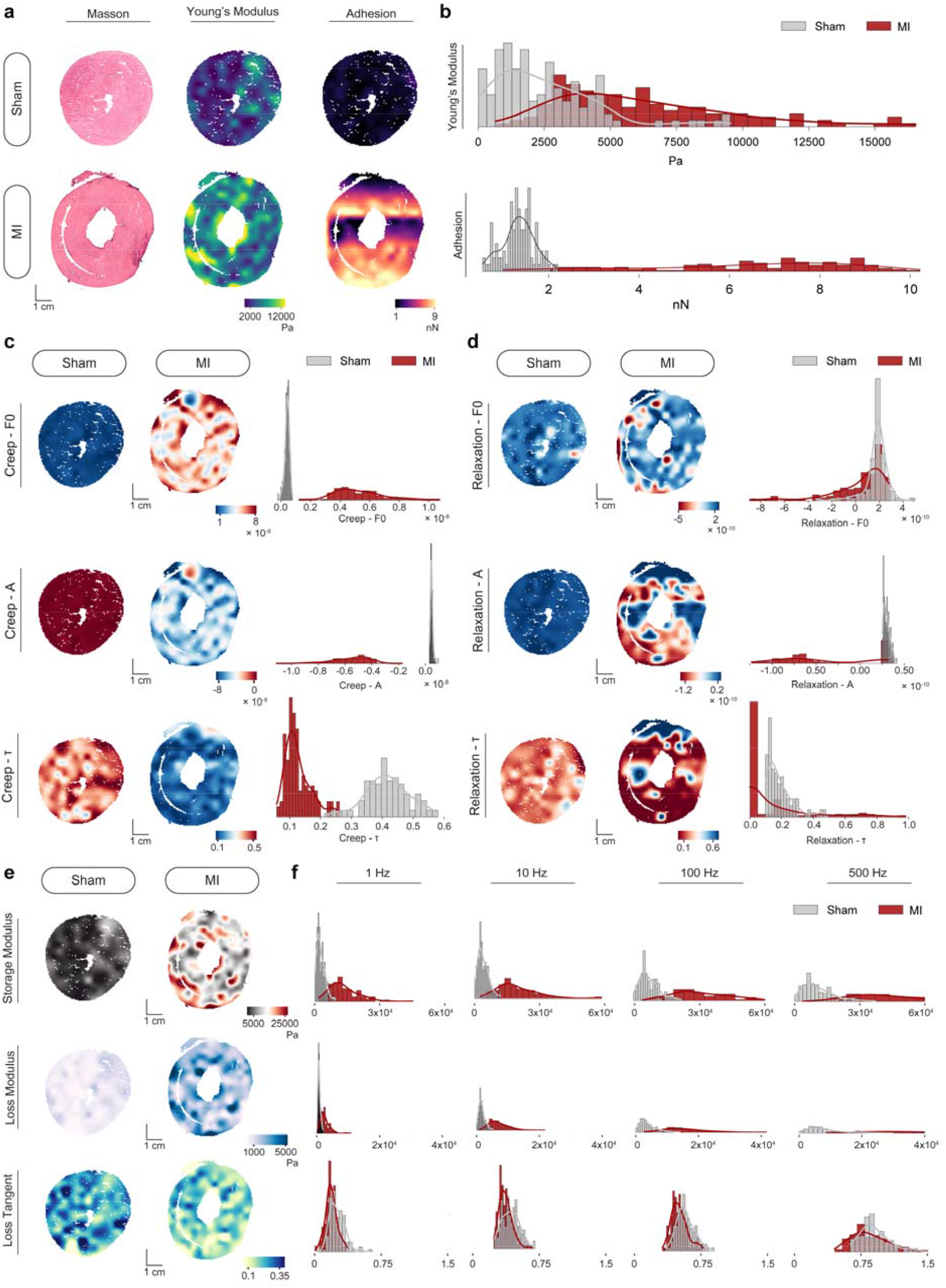
Spatial mechanomic atlases reveal tissue-scale microrheological organization. (a) Representative tissue sections with Masson’s trichrome staining and spatial maps of Young’s modulus and adhesion in control and myocardial infarction (MI) cross-section samples. (b) Distributional comparison of adhesion and Young’s modulus. (c) Spatial maps and distributions of creep-derived parameters. (d) Spatial maps and distributions of relaxation-derived parameters. (e) Frequency-dependent storage modulus, loss modulus, and loss tangent reconstructed from oscillatory microrheology. (f) Population-level comparisons across frequencies highlight condition-dependent mechanical spectra.

MI tissue exhibited a markedly stiffer and mechanically remodeled phenotype across all measured parameters (Fig. 3b–f). Young’s modulus was 2.45-fold higher in MI versus Sham (MI: 6,208 ± 3,317 Pa, median 5,420 Pa; Sham: 2,534 ± 2,001 Pa, median 2,119 Pa; p_adj_ = 1.80 × 10□^23^, rank-biserial r = −0.712), with both distributions exhibiting right-skewed, approximately lognormal character typical of soft biological tissues (Fig. 3b). Adhesion force was 4.7-fold higher in MI tissue (MI: 6.32 ± 2.34 nN; Sham: 1.33 ± 0.35 nN; p_adj_ = 8.93 × 10□□^1^, r = −0.957), indicating substantially increased cell–surface interaction in infarcted tissue.

Viscoelastic time-domain parameters revealed divergent relaxation kinetics between conditions (Fig. 3c– d). MI tissue exhibited a 3.3-fold shorter creep time constant (τ_creep_ = 0.127 ± 0.043 s in MI versus 0.415 ± 0.072 s in Sham; p_adj_ = 3.98 × 10□□^3^, r = 0.998), indicating faster viscous flow in diseased tissue. Stress relaxation τ was also reduced in MI (0.117 ± 0.210 s versus 0.187 ± 0.083 s; p_adj_ = 2.48 × 10□^12^, r = 0.506). These shorter relaxation times, coupled with increased stiffness, suggest that MI tissue has undergone structural remodeling—likely collagen deposition and cross-linking—that constrains the viscous dissipation characteristic of the native extracellular matrix.

Frequency-dependent dynamic mechanical analysis corroborated and extended the quasi-static findings (Fig. 3e–f). Storage modulus E′ was elevated 2.6- to 4.5-fold across all frequencies in MI tissue (e.g., E′ at 1 Hz: +352.2%, p_adj_ = 3.00 × 10□□□; E′ at 500 Hz: +263.6%, p_adj_ = 6.58 × 10□^3^□). Loss modulus E′′ followed a similar trend with 2.5- to 3.5-fold increases. Loss tangent (tan δ) was modestly but significantly decreased in MI at all frequencies (e.g., tan δ at 1 Hz: −26.7%, p_adj_ = 8.39 × 10□^11^), indicating a shift toward a more elastic, less dissipative mechanical response. All 20 mechanical parameters reached significance after Benjamini–Hochberg FDR correction (q < 0.05), confirming a global mechanical remodeling phenotype in post-infarction cardiac tissue. Spatial reconstruction of these parameters revealed coherent mechanical domains within each tissue section that correspond to histologically identifiable regions, establishing that spatial mechanomics captures biologically meaningful tissue architecture.

### Mechanomic feature representations enable spatial domain discovery and phenotypic discrimination

A central goal of the spatial mechanomics framework is to represent each tissue niche as a multi-dimensional mechanical feature vector suitable for unsupervised analysis, analogous to gene expression profiles in spatial transcriptomics. To assess whether the 20-parameter mechanomic representation captures biologically meaningful structure, we performed dimensionality reduction, clustering, correlation analysis, and effect size quantification across MI and Sham conditions (Fig. 4).

**Fig. 4.**
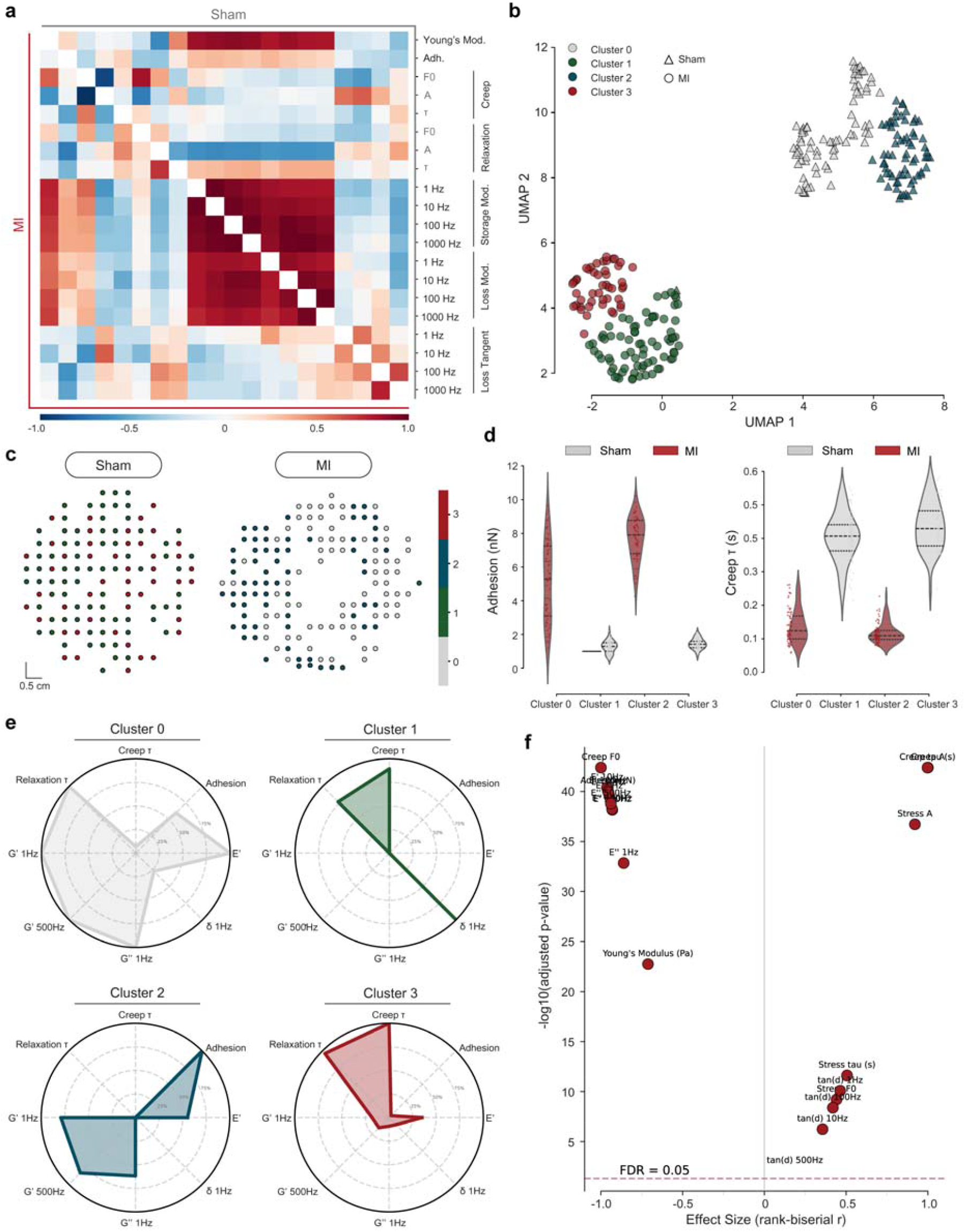
Mechanomic feature representation enables spatial domain discovery and phenotypic discrimination. (a) Mechanomic feature matrices reveal correlations among mechanical parameters across samples. (b) UMAP of mechanomic features reveals distinct mechanical states and condition-dependent separation. (c) Spatial projection of mechanomic clusters identifies tissue domains defined by mechanical phenotype. (d) Cluster-specific distributions of adhesion and creep time across samples. (e) Per-cluster mechanical fingerprints characterize distinct mechanical phenotypes. (f) Effect size analysis identifies features contributing to condition-dependent mechanical differences.

Pearson correlation analysis of the mechanomic feature matrices revealed both conserved and condition-specific inter-parameter coupling structures (Fig. 4a). The strongest conserved correlation was between Young’s modulus and storage modulus at 1 Hz (r = 0.769 in MI; r = 0.856 in Sham), confirming the expected physical coupling between quasi-static and low-frequency dynamic stiffness. However, the mechanoadhesive coupling between Young’s modulus and adhesion differed markedly: MI tissue showed negligible correlation (r = −0.038) while tissue in Sham group exhibited a modest positive association (r = 0.205). This decorrelation suggests that disease-associated stiffening and adhesion remodeling are driven by independent pathological processes—for example, collagen deposition driving stiffness and altered surface glycocalyx or integrin expression modulating adhesion. The coupling between Young’s modulus and creep time constant was near-zero in both conditions (MI: r = −0.048; Sham: r = −0.047), indicating that elastic stiffness and viscous relaxation kinetics are independently regulated regardless of disease state.

Uniform Manifold Approximation and Projection (UMAP) applied to the standardized 20-feature mechanomic space produced a well-separated two-dimensional embedding in which MI and Sham niches occupied distinct manifold regions (Fig. 4b). Principal component analysis confirmed that the dominant axis of mechanical variation aligns with disease state, with PC1 alone capturing 59.7% of total variance and PC1–PC3 explaining 79.4% cumulatively. K-means clustering (k = 4) on the UMAP embedding yielded a silhouette score of 0.455, with four clusters that partitioned almost exclusively by sample identity: Cluster 0 (n = 77, 100% MI, mean E = 7,427 Pa), Cluster 1 (n = 76, 98.7% Sham, mean E = 1,765 Pa), Cluster 2 (n = 67, 100% MI, mean E = 4,891 Pa), and Cluster 3 (n = 47, 100% Sham, mean E = 3,737 Pa). The near-complete sample segregation in unsupervised feature space demonstrates that the mechanomic representation is sufficient to discriminate disease states without supervised labels.

Critically, the identification of two subclusters per condition reveals intra-tissue mechanical heterogeneity: MI tissue comprises a stiff subpopulation (Cluster 0) and a softer subpopulation (Cluster 2), while Sham tissue partitions into compliant (Cluster 1) and relatively stiffer (Cluster 3) domains. Spatial projection of cluster assignments onto physical tissue coordinates revealed that these mechanomic clusters form spatially coherent domains rather than random interspersions (Fig. 4c–e), demonstrating that mechanically distinct niches are organized into contiguous tissue zones—consistent with the spatial organization of fibrotic remodeling in the MI condition and native tissue architecture in the Sham condition.

A volcano-style effect size analysis summarized the comprehensive statistical comparison across all 20 mechanomic parameters (Fig. 4f). All parameters reached FDR significance (q < 0.05), with the majority exhibiting large effect sizes (|r| > 0.4). The most extreme effects were observed for creep model parameters (creep F□: r = −1.000; creep A: r = 1.000; p_adj_ = 3.94 × 10□□^3^), indicating complete distributional separation between conditions. Dynamic moduli formed a cluster of large-effect, highly significant parameters, while loss tangent occupied a distinct region with moderate positive effect sizes (r = 0.19–0.46). This multi-parametric pattern confirms that post-infarction mechanical remodeling is not limited to a single property class but manifests as a coordinated, holistic transformation encompassing elasticity, adhesion, viscoelastic relaxation, and frequency-dependent dynamic behavior, supporting the concept of a comprehensive mechanical phenotype that spatial mechanomics is uniquely positioned to characterize.

## Discussion

We have introduced spatial mechanomics as a quantitative framework for tissue-scale biomechanical profiling, implemented through a multi-protocol BioAFM measurement strategy and the MechScape computational platform. By treating each spatial position as a multi-parametric mechanical niche— analogous to a pixel in spatial transcriptomics—our approach transforms tissue mechanics from isolated, single-parameter measurements into a systematic, high-dimensional omics layer amenable to the analytical toolkit of modern data science.

Several aspects of this framework represent methodological advances. First, the multi-protocol force spectroscopy design captures time-domain and frequency-domain viscoelastic behavior within a single measurement cycle, avoiding the registration errors and throughput limitations of sequential single-mode measurements. Second, the assembly of extracted parameters into per-niche feature vectors and their spatial reconstruction creates a data structure that is formally analogous to spatial omics feature matrices. This representation enables direct application of unsupervised learning methods—dimensionality reduction, clustering, correlation analysis—that have proven transformative in spatial genomics. Our UMAP analysis demonstrates that the 20-parameter mechanomic representation carries sufficient information to discriminate disease states without supervised labels (silhouette score 0.455) and, moreover, to resolve intra-condition heterogeneity through the identification of mechanically distinct subpopulations within each tissue type. Third, the biological findings enabled by spatial mechanomics offer insights not accessible through conventional mechanical measurements. The observation that adhesion and stiffness are decorrelated in MI tissue (r = −0.038) but modestly coupled in Sham tissue (r = 0.205) suggests that these properties are independently regulated in disease, potentially reflecting distinct contributions of collagen deposition (stiffening) and altered surface molecular composition (adhesion remodeling). Similarly, the frequency-dependent reduction in loss tangent across all frequencies in MI tissue reveals a shift toward more elastic, less dissipative behavior—a signature of increased cross-linking and reduced matrix hydration that cannot be captured by quasi-static elasticity measurements alone. The spatial coherence of mechanomic clusters, which form contiguous tissue domains rather than random patterns, provides evidence that mechanical remodeling follows biologically organized spatial programs consistent with the progression of fibrotic scarring from the infarct core.

Several limitations warrant discussion. The current throughput of BioAFM-based spatial sampling (approximately 15.5 s per niche) limits the number of spatial positions that can be practically measured per tissue section, resulting in coarser spatial resolution compared with imaging-based omics modalities. Advances in high-speed AFM and parallelized cantilever arrays may alleviate this constraint. The constitutive models employed, while physically interpretable, assume linear viscoelasticity under small deformations; nonlinear or strain-stiffening behavior, which is relevant in many biological contexts, would require extension of the model library. Additionally, the current validation focuses on cardiac tissue in a single disease model; demonstration across diverse tissue types and pathologies will be necessary to establish generalizability.

In summary, spatial mechanomics establishes mechanics as a quantitative, spatially resolved omics modality. By enabling systematic measurement, quantification, spatial mapping, and multimodal integration of tissue mechanical properties, this framework positions the mechanical microenvironment as a first-class variable in tissue biology and complements molecular profiling with the physical context in which molecular programs operate.

## Methods

### Animal model and tissue preparation

All animal experiments were performed in accordance with institutional guidelines and approved by the local ethics committee. Myocardial infarction was induced in adult C57BL/6J mice by permanent ligation of the left anterior descending coronary artery. Sham-operated animals underwent identical surgical procedures without ligation. Hearts were harvested at 7 days post-surgery, embedded in optimal cutting temperature (OCT) compound, flash-frozen in liquid nitrogen–cooled isopentane, and cryosectioned at 10 µm thickness. Serial sections were collected on glass slides for AFM-based mechanomics and Masson’s trichrome histological staining.

### BioAFM spatial force spectroscopy

AFM microrheology measurements were performed using a JPK NanoWizard V (Bruker/JPK) in liquid. A silicon nitride MLCT-E cantilever (triangular geometry) was used with a calibrated spring constant k = 63.23 mN/m. The tip was modeled as a sphere with radius R = 6 µm. A 10-segment programmable force protocol was executed at each grid position, totaling approximately 15.5 s per curve across all segments. Segments comprised: (0) z-extend approach to a force trigger setpoint of 1.29 V (85 ms, 174 points); (1) constant-height creep hold (2.0 s, 4,100 points at 2,050 Hz); (2) 10-nm z-retract recovery step (5 ms); (3) force-triggered retraction to 0.48 V setpoint; (4) constant-force clamp with feedback control (i-gain = 50, p-gain = 0.001; 2.0 s, 4,096 points at 2,048 Hz); (5–8) sinusoidal z-modulation at 1, 10, 100, and 500 Hz with 4 nm commanded amplitude; and (9) full z-retract of 5 µm (2.5 s). Raw data were recorded as signed 32-bit big-endian integers for height, vDeflection, and capacitive measuredHeight channels and converted to physical units using the JPK calibration chain.

### Creep and stress relaxation model fitting

Force relaxation during the constant-height creep segment was fitted with four constitutive models by nonlinear least-squares minimization: (i) standard linear solid (SLS/Zener): F(t) = F∞ + (F□ − F∞)exp(−t/τ), 3 parameters; (ii) two-arm generalized Maxwell: F(t) = F∞ + A□ exp(−t/τ□) + A□ exp(−t/τ□), 5 parameters; (iii) KWW stretched exponential: F(t) = F∞ + (F□ − F∞)exp[−(t/τ)□], 4 parameters; and (iv) power-law: F(t) = F∞ + (F□ − F∞)(1 + t/t□)□□, 4 parameters. Model selection was performed via the corrected Akaike Information Criterion: AICc = n·ln(RSS/n) + 2k + 2k(k + 1)/(n − k − 1), where n is the number of data points, k is the number of parameters, and RSS is the residual sum of squares. Instantaneous and equilibrium moduli (E□, E∞) were derived from the fitted F□ and F∞ using Hertz contact mechanics for a spherical indenter: F = (4/3)E*√R·δ^3^□

Under constant-force feedback, the z-height creep compliance was fitted with analogous models: (i) SLS: h(t) = h□ + Δh∞(1 − exp(−t/τ)); (ii) two-arm generalized Maxwell: h(t) = h□ + A□(1 − exp(−t/τ□)) + A□(1 − exp(−t/τ□)); (iii) KWW: h(t) = h□ + Δh∞(1 − exp[−(t/τ)□]); and (iv) power-law: h(t) = h□ + At□. For spatial mapping (Figs. 3–4), a single-exponential SLS model was used uniformly across all niches to enable direct parameter comparison across tissue positions. Model selection was again by AICc.

### Oscillatory microrheology

Sinusoidal z-modulation was applied at f = 1, 10, 100, and 500 Hz with a commanded amplitude of 4 nm. Force and capacitive measured-height signals were Fourier-transformed (Hanning-windowed FFT) and the amplitude and phase at the drive frequency were extracted. The complex shear modulus was computed as G*(w) = Fω / (2a·δω), where a = √(R·δ□) is the contact radius and the subscript ω denotes oscillatory amplitudes. Storage and loss moduli were decomposed as G′(ω) = |G*|cosφ and G′′(ω) = |G*|sinφ, with φ the phase lag of force behind indentation. The frequency-dependent spectra were simultaneously fitted with four rheological models (SLS, two-arm generalized Maxwell, fractional power-law, and fractional Kelvin–Voigt), using weighted residuals and AICc for model selection.

### Comparative distribution analysis

Raw mechanomics data were loaded from YAML descriptors using the MechScape data framework. Data cleansing included removal of entries with missing spatial coordinates, replacement of infinite values with NaN, and IQR-based outlier removal (1.5× IQR) applied independently per sample for Young’s modulus, adhesion, creep τ, and stress relaxation τ. For each of the 20 mechanical parameters, kernel density estimation (KDE; Gaussian kernel, Scott’s rule bandwidth) was overlaid on normalized histograms. Two-sided Mann–Whitney U tests were employed for pairwise comparison between MI and Sham groups. Multiple testing correction was applied across all 20 parameters using the Benjamini–Hochberg FDR procedure at q < 0.05. Effect sizes were quantified using the rank-biserial correlation coefficient: r = 1 − 2U/(n□n□), where U is the Mann–Whitney statistic and n□, n□ are the respective sample sizes.

### Dimensionality reduction and clustering

UMAP was used for nonlinear dimensionality reduction of the 20-parameter mechanomic feature space. All parameters were standardized (zero mean, unit variance) prior to analysis. PCA was performed on the standardized data to assess linear variance structure. UMAP was applied with n□neighbors = 15 and min□dist = 0.1 (Euclidean metric), projecting each niche into a two-dimensional embedding. K-means clustering (k = 4) was performed on the UMAP embedding to identify mechanically distinct subpopulations. Cluster quality was assessed via the silhouette score. Spatial cluster maps were generated by plotting UMAP cluster assignments at physical AFM coordinates. Pairwise inter-cluster significance was assessed using Kruskal–Wallis tests followed by pairwise Mann–Whitney U tests with FDR correction.

### Correlation and effect size analysis

Pairwise Pearson correlation matrices were computed independently for MI and Sham samples across all 20 mechanical parameters. Upper-triangle correlation heatmaps were generated for visual comparison of correlation architecture. A volcano-style visualization was constructed by plotting the rank-biserial effect size r on the x-axis against −log□□(p_adj_) on the y-axis for each parameter. Parameters reaching FDR significance (q < 0.05) were highlighted and annotated.

### Software implementation

MechScape was implemented in Python (≥3.9) as an open-source napari plugin distributed under the MIT licence. The software builds on the napari image viewer (≥0.4.19) and its npe2 plugin engine, with core numerical routines provided by NumPy, SciPy, pandas, scikit-image, and scikit-learn. MechScape adopts a YAML-centric data model in which each omics layer—AFM mechanomics, spatial transcriptomics (10× Visium), spatial metabolomics (MALDI/DESI), or multiplex immunofluorescence—is described by a portable YAML descriptor. MechScape operates both as a standalone command-line viewer and as a native napari dock-widget plugin.

## Code and Data Availability

MechScape source code is available at https://github.com/wc-xie/MechScape. Documentation is hosted at https://mechscape-docs.readthedocs.io. All raw AFM force curve data and processed mechanomic feature matrices used in this study will be deposited in a public repository upon formally publication.

## Acknowledgements

We thank the members of the laboratory for helpful discussions and critical reading of the manuscript. This work was supported by National Science Foundation for Distinguished Young Scholars of China (82125019).

## References

1. Discher, D. E., Janmey, P. & Wang, Y. Tissue cells feel and respond to the stiffness of their substrate. Science 310, 1139–1143 (2005).

2. Engler, A. J., Sen, S., Sweeney, H. L. & Discher, D. E. Matrix elasticity directs stem cell lineage specification. Cell 126, 677–689 (2006).

3. Humphrey, J. D., Dufresne, E. R. & Schwartz, M. A. Mechanotransduction and extracellular matrix homeostasis. Nat. Rev. Mol. Cell Biol. 15, 802–812 (2014).

4. Vogel, V. & Sheetz, M. Local force and geometry sensing regulate cell functions. Nat. Rev. Mol. Cell Biol. 7, 265–275 (2006).

5. Iskratsch, T., Wolfenson, H. & Sheetz, M. P. Appreciating force and shape—the rise of mechanotransduction in cell biology. Nat. Rev. Mol. Cell Biol. 15, 825–833 (2014).

6. Hinz, B. et al. Recent developments in myofibroblast biology: paradigms for connective tissue remodeling. Am. J. Pathol. 180, 1340–1355 (2012).

7. Plodinec, M. et al. The nanomechanical signature of breast cancer. Nat. Nanotechnol. 7, 757–765 (2012).

8. Berry, M. F. et al. Mesenchymal stem cell injection after myocardial infarction improves myocardial compliance. Am. J. Physiol. Heart Circ. Physiol. 290, H2196–H2203 (2006).

9. Fung, Y. C. Biomechanics: Mechanical Properties of Living Tissues (Springer, 1993).

10. Radmacher, M. Studying the mechanics of cellular processes by atomic force microscopy. Methods Cell Biol. 83, 347–372 (2007).

11. Krieg, M. et al. Atomic force microscopy-based mechanobiology. Nat. Rev. Phys. 1, 41–57 (2019).

12. Ståhl, P. L. et al. Visualization and analysis of gene expression in tissue sections by spatial transcriptomics. Science 353, 78–82 (2016).

13. Asp, M., Bergenstrahle, J. & Lundeberg, J. Spatially resolved transcriptomes—next generation tools for tissue exploration. BioEssays 42, e1900221 (2020).

14. Rao, A., Barkley, D., França, G. S. & Yanai, I. Exploring tissue architecture using spatial transcriptomics. Nature 596, 211–220 (2021).

